# HlyF, an underestimated virulence factor of uropathogenic *Escherichia coli*

**DOI:** 10.1101/2023.04.27.538512

**Authors:** Camille V. Chagneau, Delphine Payros, Audrey Goman, Cécile Goursat, Laure David, Miki Okuno, Pierre-Jean Bordignon, Carine Séguy, Clémence Massip, Priscilla Branchu, Yoshitoshi Ogura, Jean-Philippe Nougayrède, Marc Marenda, Eric Oswald

**Affiliations:** Digestive Health Research Institute (IRSD), INSERM, Université de Toulouse, INRAE, ENVT, UPS, Toulouse, France; CHU Toulouse, Hôpital Purpan, Service de Bactériologie-Hygiène, Toulouse, France; Division of Microbiology, Department of Infectious Medicine, Kurume University School of Medicine, Kurume, Japan; Faculty of Science, University of Melbourne, Melbourne, Australia

**Author notes:** corresponding author, Pr Eric Oswald, Institut de recherche en santé digestive, CHU Purpan – Place Baylac, CS 60039, 31024 TOULOUSE CEDEX 3, France.

**Keywords:** uropathogenic *E. coli*, urinary tract infections, HlyF, ColV plasmids

## Abstract

Urinary tract infections (UTIs) are predominantly caused by uropathogenic *Escherichia coli* (UPEC). By analysing a representative collection of UPEC strains from community-acquired infections, we showed that 20 % of these strains had the ability to produce the protein HlyF. These *hlyF*+ UPEC strains were the most virulent, mostly responsible for pyelonephritis, often with bloodstream infections. Using a mouse model of UTI, we showed that HlyF was associated with the ability of UPEC to develop a urosepsis, with the presence of bacteria in the spleen and an exacerbated inflammatory response. In contrast to archetypical UPEC strains, *hlyF*+ UPEC strains are not restricted to phylogroup B2 and harbor a specific repertoire of virulence factors reflecting the fact that HlyF is encoded by conjugative ColV-like plasmids. These plasmids also carry antimicrobial resistance genes, which may facilitate their selection and spreading amongst people receiving antimicrobial therapy. Overall, our data suggest that HlyF is a virulence factor in UPEC and spreading of ColV-like plasmids encoding *hlyF* warrants further investigation.

## Introduction

Urinary tract infections (UTIs) are one of the most common infections worldwide [1]. UTIs are associated with a decrease in the quality of life of patients and a significant clinical and economic burden [2]. In both community and hospital settings, UTIs pose a threat to public health. They are the most common outpatient infections and at least half of adult women will have one UTI or more in their lifetime [3]. Uropathogenic *E. coli* (UPEC) are responsible for more than three quarters of community-acquired UTI, and about half of nosocomial infections [4]. The majority of these infections are benign, but their management can be complicated by frequent recurrences and the emergence of antibiotic resistance, leading to therapeutic impasses. In more severe cases, complications such as kidney damage in young children or the onset of sepsis may arise. A large proportion of sepsis originates from the urinary tract (accounting for 20-30%, *i.e*. 2 to 9 million cases per year) and urosepsis may progress to septic shock with significant mortality and morbidity [5]. The mortality from urosepsis is estimated to be more than 1.5 million deaths per year, making it a major public health threat.

The pathogenicity of UPEC involves a variety of factors [6] such as specific adhesins, toxins or iron uptake systems [7,8]. The *hlyF* gene encodes a protein that was previously thought to be an haemolysin (haemolysin F) [9]. Recent work has shown that HlyF is in fact a cytoplasmic enzyme that increases the formation of outer membrane vesicles (OMVs) allowing the release of the *bona fide* haemolysin E (ClyA) responsible for the previously observed haemolytic phenotype [10]. HlyF-induced OMVs not only act as cargos for toxins, but also have the ability to block autophagic flux in eucaryotic cells and to exacerbate the activation of the inflammasome through the non-canonical pathway [10,11].

So far the *hlyF* gene was shown to be associated with the virulence of avian pathogenic *E. coli* (APEC) and neonatal meningitis-causing *E. coli* (NMEC) [10,12–14]. In this study, we observed that *hlyF+* UPEC were isolated from the most severe cases of human UTI. The ColV plasmids carrying *hlyF* are widely spread among UPEC strains and encode several virulence factors as well as antimicrobial resistances which can favour their dissemination. In a mouse model of UTI, we have shown that HlyF promoted pyelonephritis and consecutive bloodstream infection.

## Materials and methods

### Collection of clinical strains

We collected 225 *E. coli* strains from prescribed urine cultures of 223 patients attending the Adult Emergency Department of Toulouse University Hospital, France, between July and October 2017, corresponding to community-acquired UTIs as previously described [15]. Patients at risk of misdiagnosis due to age (> 75 years) or comorbidities, and patients with urinary catheters were excluded. In accordance with French regulations on the analysis of observational databases, no specific informed consent was required for the collection of clinical *E. coli*. Data were analysed anonymously.

### Bacterial strains

UPEC strain ECC166 was isolated from a 23-year-old woman without comorbidities suffering from pyelonephritis. Whole genome sequence of ECC166 indicates it is of serotype O1:H7, phylogroup B2 and sequence type ST95. In addition to the ability to produce HlyF, ECC166 exhibits a wide array of virulence factors such as multiple iron acquisition systems (locus *iro, iucABCD, fyuA, sitABCD*), Vat toxin and PapGII adhesin. We constructed a deletion mutant of *hlyF* in UPEC strain ECC166 as previously described [16]. The Δ*hlyF* mutant was constructed using primers PB3-mut-hlyF-F (5’- TAAGATAATTTATTTTTATAATGATCACATGAAAACAAAAGAGGTTAGATgtg taggctggagctgcttcg-3’) and PB4-mut-hlyF-R (5’- TTTATATATTATGAGTGCAACACCAACAATAATTCTGATTATGATAAATAcata tgaatatcctccttagt-3’). This mutant was complemented with plasmids expressing a wild-type form of HlyF (referred to as HlyF) or a mutant in the catalytic domain (referred to as SDM) [10].

### Mouse model of UTI

#### Ethic statement

All the experimental procedures were carried in accordance with the European directives for the care and Use of animals for Research Purposes and were validated by the local ethics committee from CREFRE US006 and by the national ethics committee (Regional Centre for the Functional Exploration and Experimental Ressources) (number 21-U1220-EO/MT-128).

#### Mouse model

Female, 6-8 weeks old, C3H/HeN mice (Janvier Labs, Le Genest Saint Isle, France) were infected twice transurethrally as previously described [17]. Briefly, bacterial strains were cultivated statically in LB and resuspended to an inoculum of 2.10^e^7 CFU in 50µL of PBS 1X. For bacterial enumeration, bladder, kidney and spleen were harvested, homogenised in FastPrep Lysing Matrix D with 800µL PBS 1X and serial dilutions were plated onto solid LB agar with adequate antibiotic.

#### Body weight and clinical scoring

Body weight was assessed before and at the end point of the experiment. The severity of the clinical signs was evaluated blindly by scoring (body temperature, coat condition, mobility of the animals and signs of pain such as grimace) (Table S1). A weight loss of more than 15 % associated with a clinical score of more than 11 led to stop the experiment and to humanely sacrifice the animal.

#### Cytokines quantification

Tissue proteins were extracted with a solution of RIPA (0.5% deoxycholic acid, 0.1% sodium dodecyl sulfate, 1% Igepal in Tris-buffered saline 1X; pH = 7.4) added with a protease inhibitor cocktail (Roche diagnostic, France Ref 11697498001). Clear lysates of spleen were processed for ELISA using commercial kits (Duoset R&D Systems, Lille, France) for Interleukin-1β (IL-1β) and Interleukin-6 (IL-6) according to the manufacturer’s instructions. Data are expressed as picograms of cytokines per milligram of tissue protein.

### Sequencing data, sequence alignments and phylogenetic analyses

#### Illumina sequencing

The Illumina NextSeq500 Mid Output platform (Integragen, Evry, France) was used to generate 2 x 150 bp paired-end reads for whole genome sequencing of the UPEC strains as already described [17].

#### Nanopore sequencing

Genomic DNAs were purified from 600 µL of overnight bacterial culture using the Wizard HWM genomic DNA prep kit (Promega) following the manufacturer’s instructions. Nanopore sequencing libraries were prepared with the rapid barcoding kit SQK-RBK004 (Oxford Nanopore) and loaded on a MinION MK-Ib device fitted with a R9.4 flowcell. Live rapid basecalling was performed with MinKnow 20.13.3 using the guppy version 4.2.2 (Oxford Nanopore). The resulting fastq reads were processed with the program guppy_barcoder (Oxford Nanopore) for quality filtering >7, demultiplexing, and barcode removal.

#### Genome analysis

Genome *de novo* assembly and analysis (search for virulence and antimicrobial resistance genes, phylogroup, plasmid typing…) were performed with the BioNumerics 7.6 software (Applied Maths), Enterobase and Center for Genomic Epidemiology (http://enterobase.warwick.ac.uk/; http://genomicepidemiology.org/). For SNP-based phylogenetic trees, core genome alignments were generated after mapping raw reads to the *E. coli* MG1655 genome using LASTAL after identifying dispersed repeats (BLASTN) and tandem repeats (trf) but without considering recombination [18]. The core genome phylogenetic tree was inferred with the Maximum-likelihood algorithm (RAxML) using Enterobase for *hlyF*+ strains [18].

Alignment, comparisons, and detection of mutations in *hlyF* locus were performed either using BioNumerics software or Clustal Omega [19] with pECOS88 as a reference (CU928146.1: 130 790… 133 302) from *E. coli* S88 serotype O45:K1:H7 isolated from neonatal meningitis and including other reference strains: *E. coli* SP15 of serotype O18:K1:H7 isolated from neonatal meningitis (AP024132.2) and *E. coli* Combat11I9 of serotype O-:H28 isolated from UTI (CP021728.1). MEGA X software was used to construct UPGMA tree of *hlyF* locus [20]. The evolutionary distances were computed using the Maximum Composite Likelihood method and are in the units of the number of base substitutions per site.

Multiple alignments for the *hlyF* and *traM-traX* sequences were performed by mafft v7.429 [21] and then phylogenetic estimations were conducted using the maximum-likelihood (ML) method implemented in RAxML-ng v.0.9.0 (--all, --bs-trees 100) [22]. The best-fit model for each phylogenetic analysis was selected by using ModelTest-NG v.0.1.6 [23]. Co-phylogenetic analysis between the ML trees was performed using the ‘cophylo’ function of the R package Phytools 0.7.80 [24]. Phylogenetic clusters were determined by RAMI [25].

Hybrid assembly and annotation of the fully assembled plasmids was performed using Unicycler and Prokka from the Galaxy interface (https://usegalaxy.org/). Plasmid linear comparison and representation was performed using Easyfig 2.2.5 software (https://mjsull.github.io/Easyfig/).

### Statistical analyses

Graphical representations of data were performed using the GraphPad Prism 8.0 (GraphPad Software, Inc, San Diego, CA). The experimental data are represented as means ± standard errors of the mean (SEM). For time-to-death experiment, the difference between the experimental groups was evaluated by the log-rank (Mantel-Cox) test. For the statistical analysis of clinical score, body weight loss and bacterial load, a Student’s t-test was used to differentiate HlyF expressing strains and strains not expressing HlyF. For cytokine analysis, statistical significance between experimental groups was determined by one-way ANOVA followed Bonferroni’s comparison test. P value of < 0.05 was considered significant. For epidemiological analysis on UPEC strains, P values were calculated using Fischer’s exact test.

## Results

### hlyF is epidemiologically associated with severe urinary tract infections in humans

We investigated the presence of *hlyF* in a representative collection of 225 sequenced UPEC strains from community-acquired UTIs [15]. This collection was shown to be representative of the commonly described UPEC collections, for classically described virulence factors, phylogeny and for antibiotic resistance profiles commonly reported in France [15,17]. We found that *hlyF* was present in 42/225 (19%) of the strains. These *hlyF*+ UPEC were present in all phylogroups (Figure S1). The majority of strains (30/42) belonged to phylogenetic group B2. However, the proportion of *hlyF*+ strains in the B2 phylogenetic group (30/155) was not higher than that of non-B2 strains (12/70; p=0.8536). Within the B2 phylogenetic group, the strains belonged to classically described sequence types in UPEC (ST), especially ST95. Most of ST95 strains carried the *hlyF* gene (25/27 ST95).

The isolation rate for *hlyF*+ UPEC strains was significantly higher in patients with pyelonephritis (26%; 27/104) compared to patients with less severe UTI, *e.g.* cystitis or asymptomatic bacteriuria (12.4%; 15/121; p=0.0104). We also observed that *hlyF*+ UPEC-infected patients had a trend towards higher prevalence of concomitant bloodstream infection (30%; 7/21; p=0.08).

### HlyF is a virulence factor of UPEC in a mouse model of UTI

To test whether HlyF plays a role during a UTI, we used a well-established mouse model of UTI based on transurethral injection of bacteria [26]. We selected an UPEC strain isolated from pyelonephritis: ECC166, of sequence type ST95 and serotype O1:K1:H7, which is representative of the majority of *hlyF*+ strains in our UPEC collection. Both ECC166 wild-type and *hlyF* mutant strains induced infection in mice. However, infection was more severe with the wild-type compared to the Δ*hlyF* isogenic mutant, with increased clinical signs (Table S1) and body weight loss (Figure S2), and ultimately higher lethality (40% versus less than 10% of mortality for the wild-type and mutant strain respectively) (Figure 1). We confirmed that this difference in virulence was due to the presence of the *hlyF* gene by complementing the mutant with a plasmid expressing HlyF (Figure S3). Mice infected with the wild-type strain had more bacteria in the kidneys than mice infected with the isogenic mutant although no difference was observed in the bladder (Figure 1C). In addition, mice infected with the *hlyF*+ complemented strain had a higher bacterial load in the spleen, confirming bloodstream infection from the urinary tract (Figure 1C). An enhanced inflammatory response with higher levels of interleukin IL-1β and IL-6 in the spleen was observed in these mice (Figure 1D). These results were confirmed in the *hlyF* mutant complemented with HlyF (Figure S3C).

**Figure 1.**
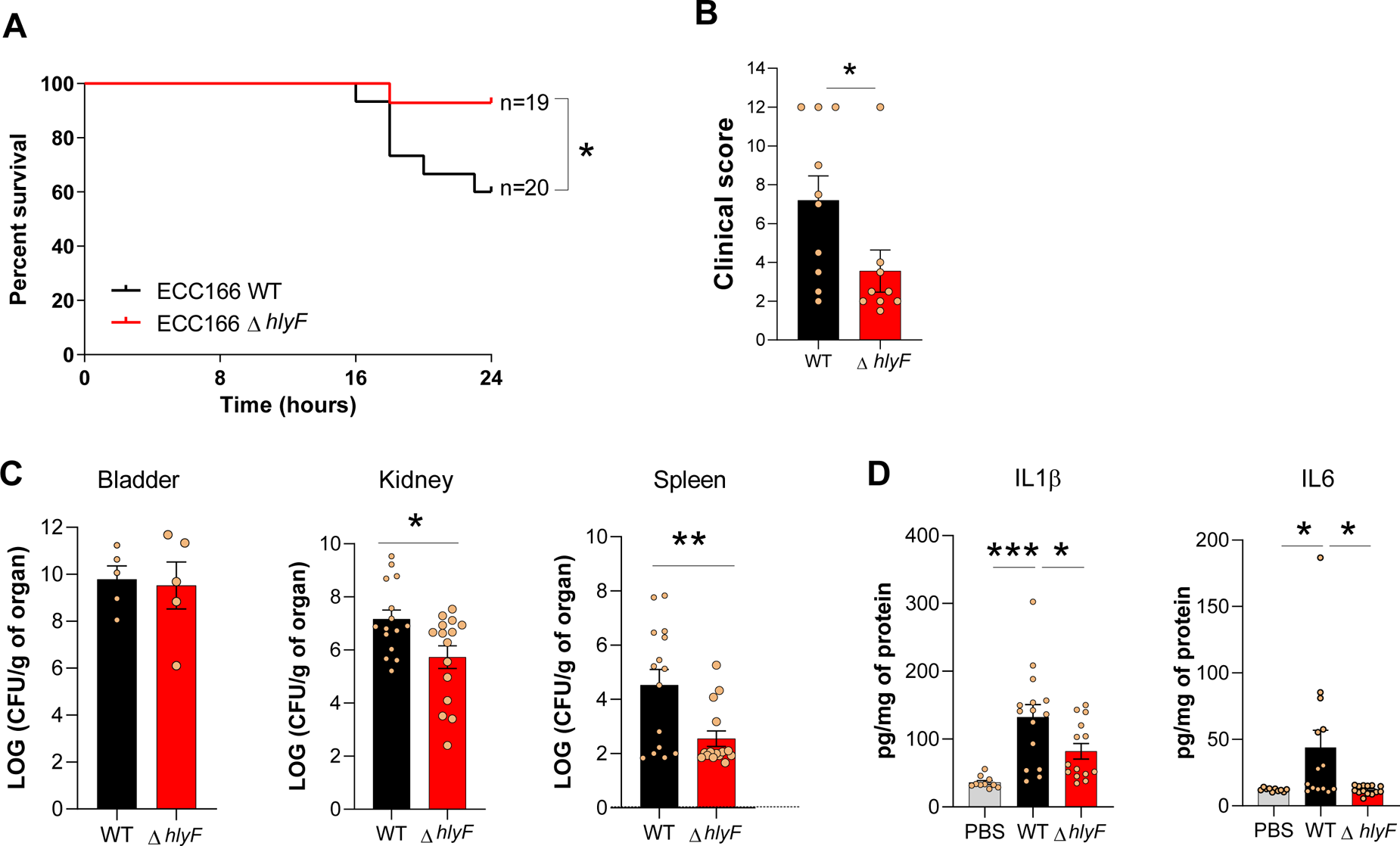
HlyF increases pathogenicity during urinary tract infection in a mouse model. C3H/HeN mice were infected transurethrally with wild-type UPEC strain ECC166 (WT) or *hlyF* isogenic mutant (Δ*hlyF*). **A.** Time to humane euthanasia (upon a body weight loss and/or clinical score reaching a predefined threshold) was monitored to build the survival curve (panel A). The results are pooled from three independent experiments, the total number of animals is shown. The difference between the experimental groups was evaluated by the log-rank (Mantel-Cox) test: * p<0.05. **B.** Clinical score according to the Table S1 at 20h+/-2h post-inoculation. Mean values ± SEM are shown, each circle represents a mouse. The presented results are pooled from two independent experiments. Student’s t-test: * p<0.05. **C.** Bacterial load in bladder, kidney and spleen (CFU/g organ) at end point. Mean values ± SEM are shown. The presented results are pooled from two independent experiments. Student’s t-test: * p<0.05; ** p<0.01 **D.** IL-1β and IL-6 in spleen (pg/mg of protein) at endpoint. Mean values ± SEM are shown. The presented results are pooled from two independent experiments. Student’s t-test: * p<0.05; *** p<0.001.

Collectively, these results indicate that the production of HlyF in UPEC strain is responsible for an increased severity of UTI in a mouse model, promotes bloodstream infection and induces a higher systemic inflammatory response.

### *HlyF+ UPEC* strains have a specific virulence signature

Using WGS of the strains collection, we compared the virulome of *hlyF*+ and of *hlyF*-UPEC strains (Table 1). Although evasion of the host response during UTI is often mediated by toxin production [27], we observed that the vast majority of *hlyF+* strains did not carry the genes for toxins classically associated with UPEC such as the alpha-haemolysin (HlyA), the Cytotoxic Necrotizing Factor 1 (CNF1), or the autotransporter protease Sat. By contrast, there was an association between *hlyF* and the adhesin PapGII, although this chromosomal allele involved in renal colonisation has been previously found almost exclusively in ST95 strains [28]. There was also a strong association with virulence genes coding for colicins, the increased serum survival type 1 plasmid variant and iron and metals scavengers and transporter systems such as aerobactin, salmochelins and the transporters encoded by *sitABCD* and *etsABC* [29,30] (Table 1). The genes encoding this arsenal of virulence factors are known to be carried by plasmids of the pColV family [29,30], which also harbour *hlyF*, raising the possibility of a common genetic origin.

**Table 1.**
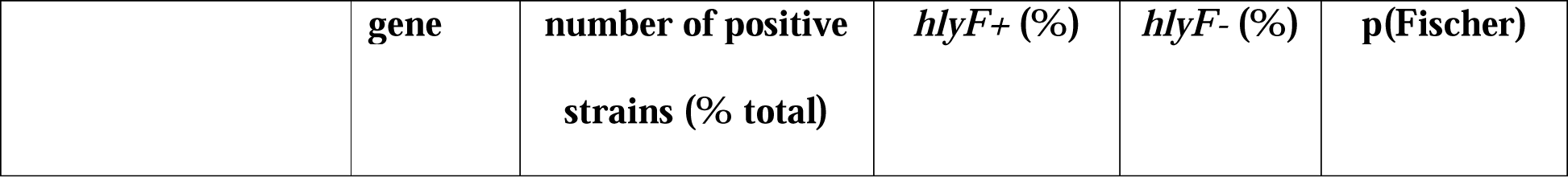
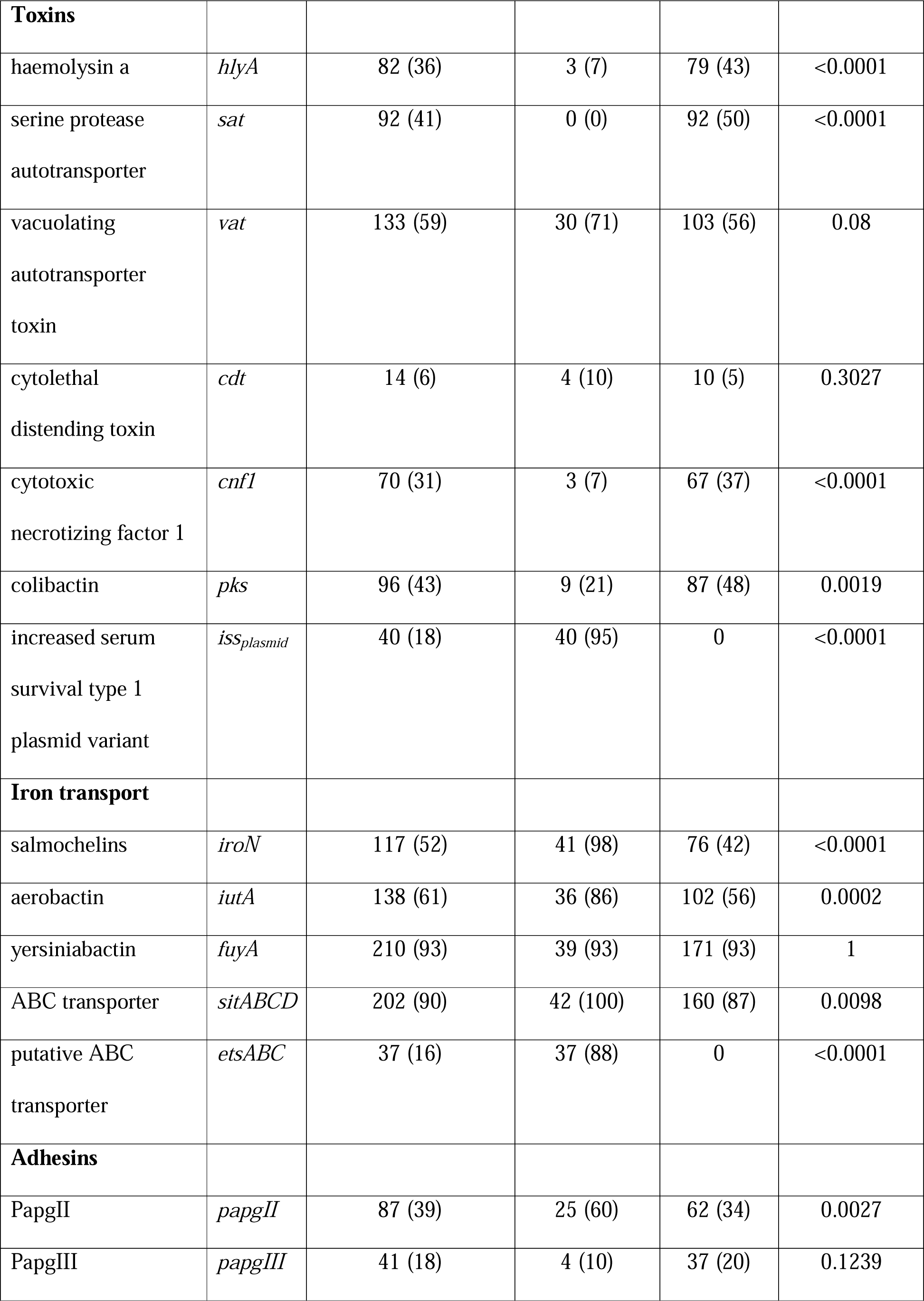
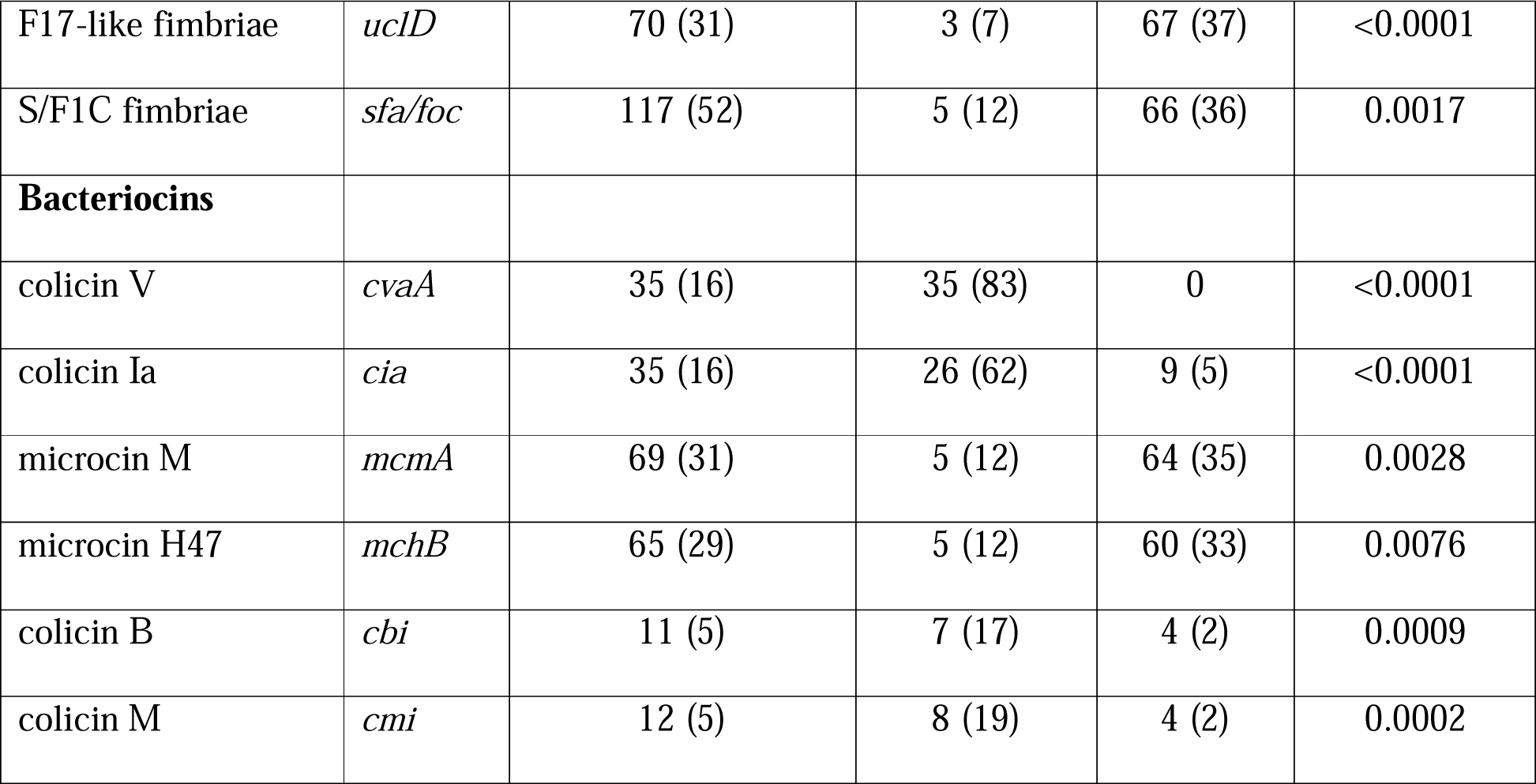
Association between HlyF and classical UPEC virulence factors. Among the strains carrying each given virulence factor, the number of *hlyF*+ and *hlyF*-strains is indicated, with the percentage among the *hlyF*+ or *hlyF-* strains in brackets. Fischer’s exact test.

### *hlyF* is carried by pColV conjugative plasmids

Assembly and annotation performed after both Nanopore and Illumina sequencing of the UPEC strain ECC166 yielded a chromosome of 4.945.664 bp and two plasmids of 115.445 bp and 1.552 bp, respectively. *hlyF* was carried by the largest plasmid belonging to pColV family (Figure S4). As previously described, *hlyF* was present in an operon found on pColV family plasmids, here referred to as *hlyF* locus, containing the *hlyF* gene and a putative *mig-14* ortholog gene (*mig-14-like*) which encodes an antimicrobial resistance factor [31–33] (Figure 2). This locus was systematically associated with *ompTp* gene, which encodes for plasmid variant of OmpT, a protease with an activity against host antimicrobial peptides [10,30,32,34]. In all UPEC strains, *hlyF* locus was highly conserved with pairwise similarities of at least 98% between two loci (Figure 2, 3 and 4). The *hlyF* gene itself showed even more similarity amongst the strains with the presence of only 11 SNP sites within the 42 strains, 6 of which generated silent mutations (Table S2). The predicted catalytic domain (YTHSK) and NAD binding domain (GATGFLG) of HlyF [10] were conserved in all strains.

**Figure 2.**
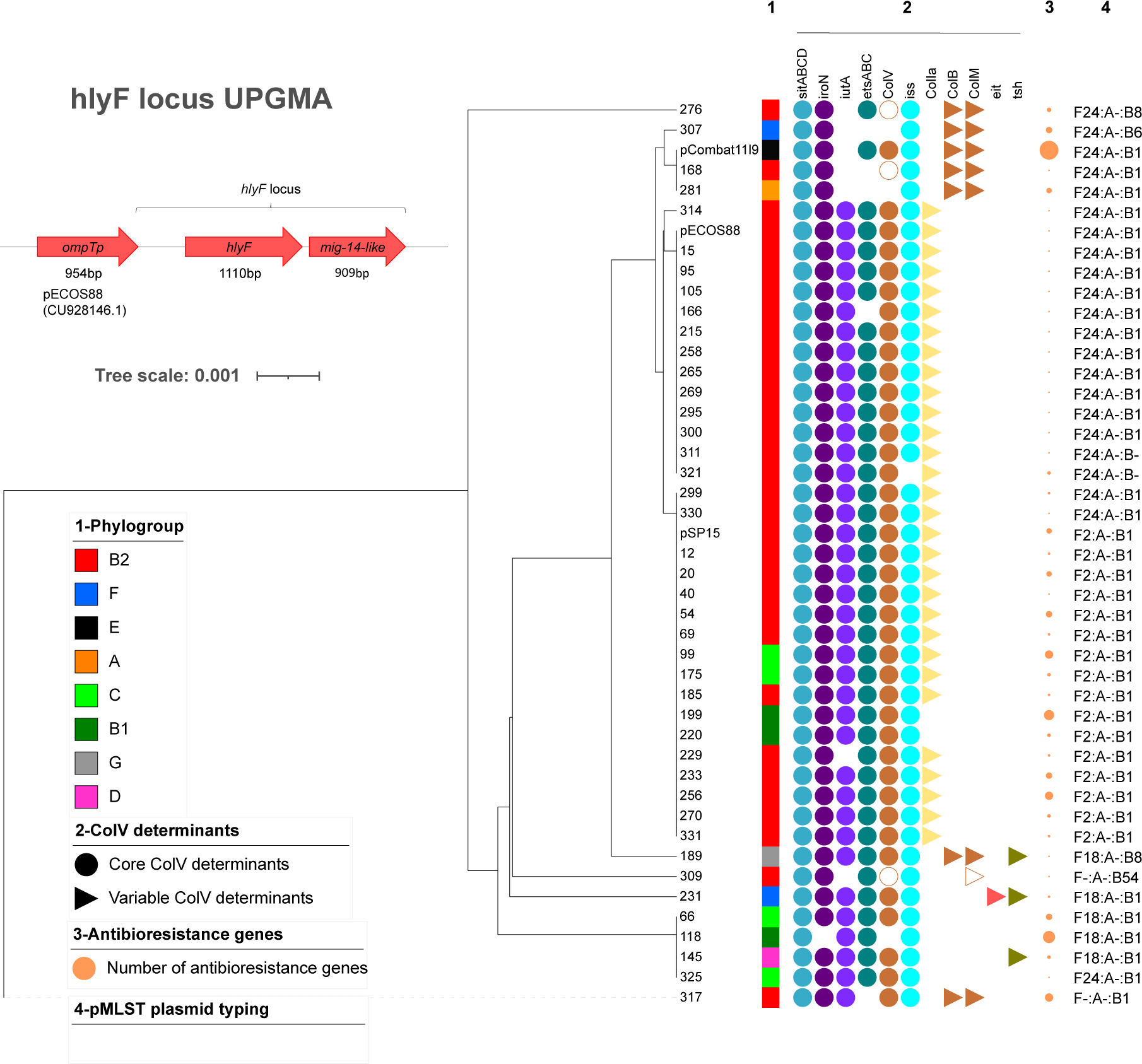
*hlyF* locus is conserved and associated with ColV plasmids determinants. The *hlyF* locus from pECOS88 used for comparison is shown together with the *ompTp* gene. Phylogenetic tree of the *hlyF* locus from UPEC reads and three fully sequenced and available genomes of plasmids from *E. coli* S88, SP15 and Combat11I9 was constructed using iTol software (https://itol.embl.de/). The phylogroup is indicated by a coloured square (column 1). The presence of classical ColV plasmid determinants is indicated by a full circle (constant determinants according to Johnson et *al*. [29]) and a full triangle (column 2). The empty version corresponds to the presence of a truncated form. The size of the circle is proportional to the number of antibiotic resistance genes found in the ResFinder search (column 3). pMLST plasmid typing is indicated column 4.

The largest plasmid of ECC166 also contained an arsenal of additional virulence factors characteristic of pColV family plasmids: colicin *colV*, aerobactin operon (*iuc/iut*), salmochelins locus *iro*, *sitABCD* metal transport system, and increased serum survival type 1 plasmid variant (*iss)* [29,30]. This plasmid also encoded a large conjugation system (*tra* genes). We confirmed its ability of transfer by conjugation, by using as a donor strain the Δ*hlyF* mutant in which a kanamycin resistance cassette was inserted in *hlyF* gene and strain *E. coli* J53 as recipient [35].

### *hlyF* is associated with two main lineages of conjugative plasmids in UPEC

The vast majority of *hlyF+* UPEC strains showed the association of *hlyF*, *ompTp*, *iutA*, *iroN, sitABCD, etsABC, iss* and *cvaA* genes which is characteristic of the conserved part of the pColV plasmids family, together with *cia,* which encodes colicin Ia [29,30] (Figure 2). We investigated whether the same plasmid was responsible for the spread of *hlyF* in the UPEC population and more broadly in the *E. coli* population. Based on the sequence of the *hlyF* locus, we identified two main groups of *hlyF*+ UPEC plasmids: one with the *hlyF* locus identical to the one of *E. coli* SP15 and one identical to the one of *E. coli* ECOS88 (Figure 2). ECOS88 and SP15 are two archetypal *E. coli* strains isolated from neonatal meningitis [36,37]. To analyse the relationship and compare the evolution of the *hlyF* locus with the structure of the plasmid carrying it, we performed a co-phylogeny analysis between the *hlyF* locus and the *traMtraX* locus, which includes the region encoding the conjugation system that constitutes the backbone of the plasmid. Interestingly, the two main groups of strains showed conserved loci and plasmid structures that have evolved in parallel (Figure S5). The gene *hlyF* is thus associated with two main lineages of plasmids in UPEC.

### *hlyF*-associated pColV plasmids are platforms for dissemination of virulence factors and antimicrobial resistance genes

In addition to the plasmids belonging to the pSP15 and pS88 lineages, more variable *hlyF* locus determinants were present in some plasmids, along with a less conserved *traMtraX* locus, associated with a diverse repertoire of pColV determinants (Figure 3 & S5), suggesting more variable plasmids. To better characterize this heterogeneity, we sequenced representative strains using the Nanopore technology and compared these plasmids with previously sequenced plasmids related to pColV (pSP15, pECOS88, pCombat11I9-2).

**Figure 3.**
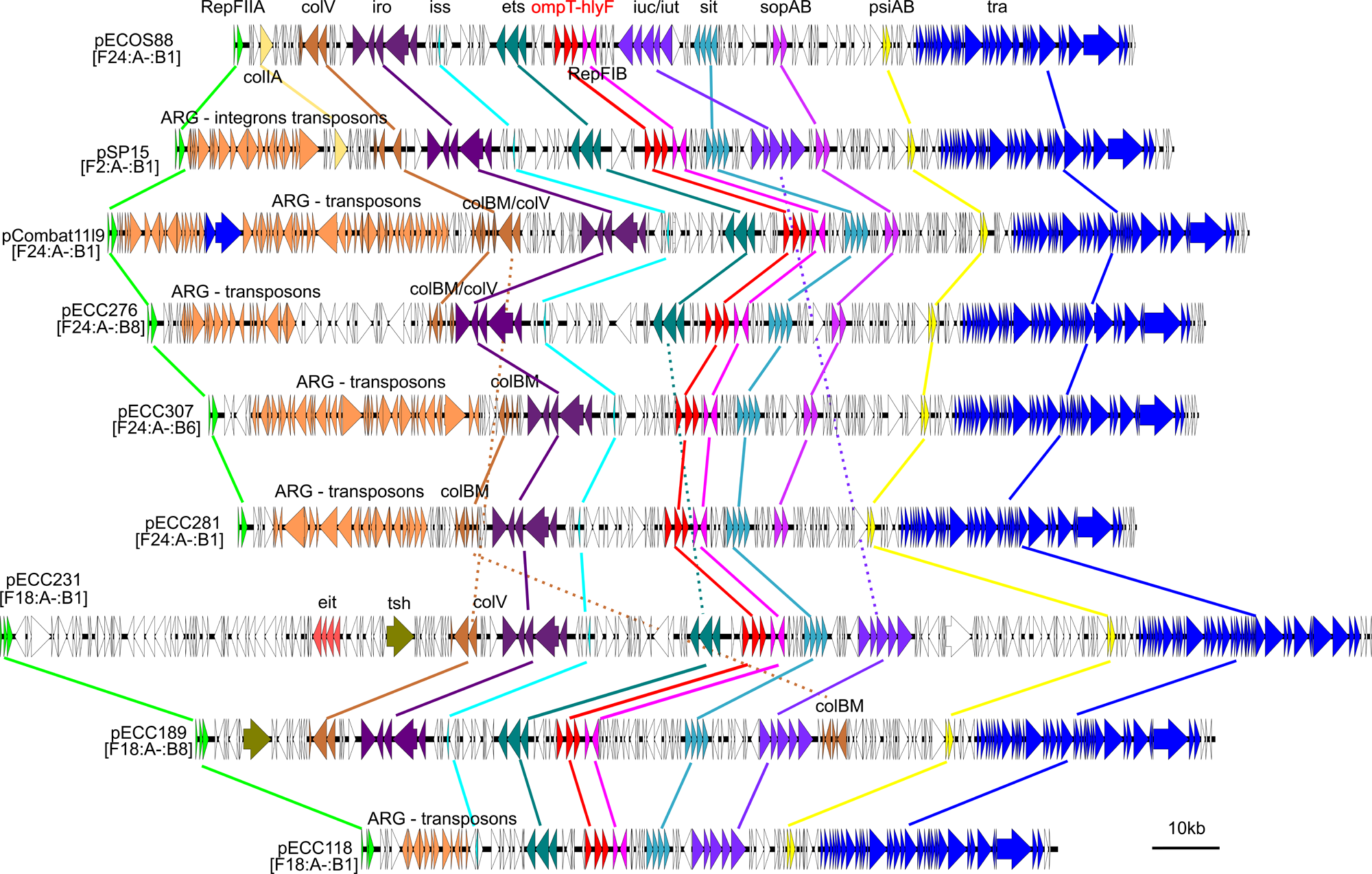
*hlyF* is carried by various mosaic plasmids combining virulence factors and antibiotic resistance genes. pMLST plasmid typing is indicated under the plasmid name. Genes of interest (virulence, plasmid replication, antimicrobial resistance genes (ARG)) are highlighted and coloured. Lines follow genes of interest throughout the figure to illustrate gene presence and overall gene conservation.

The structure of all the sequenced plasmids was conserved (Figure 3 and 4). However, many inversion sequences were present in less conserved regions, suggesting frequent IS-mediated recombinations (Figure 4). In addition, some plasmids were missing certain regions or new regions were present. These new insertions affected either (1) accessory virulence factors that are less frequently associated with pColV plasmids (*eit, tsh*), or (2) antimicrobial resistance genes carried by transposons or integrons. These results indicated that plasmids carrying *hlyF* in UPECs had a conserved scaffold, but that they were also platforms for the dissemination of various virulence and antimicrobial resistance genes. In summary, *hlyF* was carried by two main lineage of conjugative plasmids that have evolved in parallel and could promote the spread of virulence factors and antibiotic resistance genes in UPEC, a feature that may have an impact on UTI epidemiology and treatment options.

**Figure 4.**
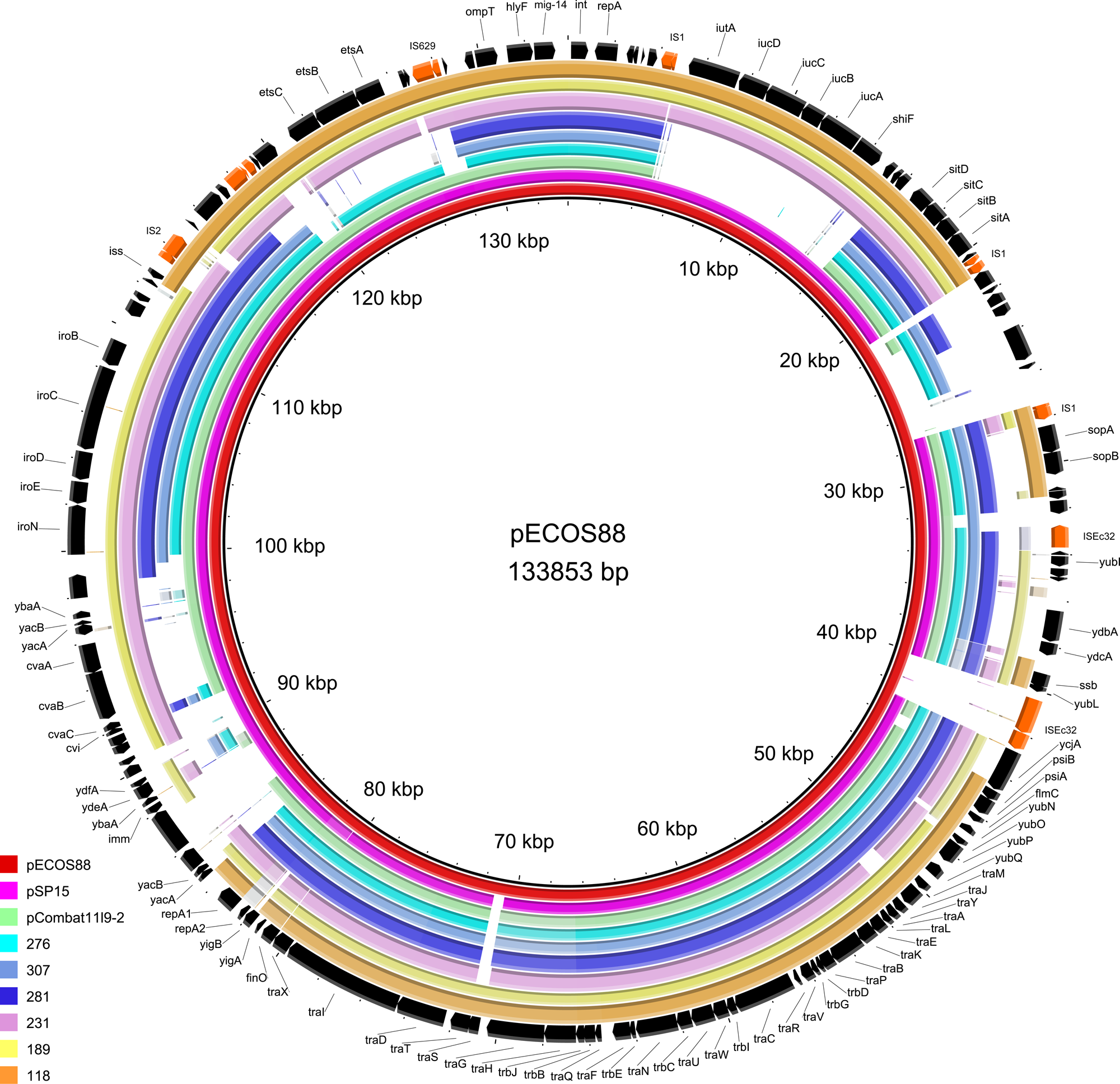
Plasmids carrying *hlyF* have a conserved scaffold but show variability. pECOS88 sequence is compared to both reference plasmids and Nanopore-sequenced plasmids from the UPEC collection using the Blast Ring Image Generator software [38]. Long tick marks on the outer and inner circumference of the ring indicate 500 kilobase pair increments and short tick marks indicate 100 kilobase pair increments. The outer black circle corresponds to ECOS88 plasmid annotation with insertion sequences highlighted in orange.

## Discussion

### Virulence plasmids and UPEC

The role of plasmids in the pathophysiology of UTI has anecdotally been studied [39]. Acquisition of the pColV plasmid may increase growth in urine and colonisation of the murine kidney [40]. The pColV family of plasmids harbour multiple virulence genes encoding iron uptake and transport functions, resistance to host response or bacteriocins that may be involved in the digestive and urinary colonisation stages of UTI. For example, both Mig-14-like and OmpTp which are highly conserved and consistently associated in *hlyF*+ UPEC strains, are involved in resistance to host antimicrobial peptides [32,33]. In particular, OmpTp is a protease that has been shown to be particularly active against antimicrobial peptides produced in the urinary tract [34]. However, independently of the other virulence factors encoded by ColV, we show in a mouse model that HlyF alone can control the severity of UTI and contribute to the occurrence of bloodstream infection.

### HlyF: an emerging virulence factor associated with severe UTI

The gene encoding HlyF has been previously described in APEC and NMEC strains [12–14] but has never been studied in UPEC, probably because *hlyF* is absent from the genome of archetypal UPEC strains such as *E. coli* CFT073, 536 or UTI89. We have observed in our representative UPEC collection that almost 20% of strains isolated from community-acquired infections carry this virulence factor. This frequency is confirmed by data from the literature based on the epidemiology of genes classically associated with *hlyF* (*ompTp*, *cvaC*…) [28,34,41–43]. *hlyF*+ strains seem especially frequent in UPEC belonging to ST95. Interestingly, in larger collections, some subgroups of ST95 strains (C and D) are much more likely to carry pColV plasmids and are becoming more prevalent in patients with UTI, suggesting an on-going dissemination [44]. These strains are mostly devoid of classical UPEC virulence genes such as *hlyA* or *cnf1* [28], which may suggest either alternative virulence mechanisms specific to HlyF or pColV determinants, or that the accumulation of the virulence genes may be too detrimental for the bacterial host and would result in strains being unable to colonise the urinary tract. We found that UPEC strains producing HlyF are epidemiologically associated with more severe UTI in humans. Interestingly, we observed a trend towards a higher frequency of bloodstream infections, in accordance with larger studies [28]. We also confirmed HlyF role *in vivo* in mouse model of UTI.

### A role of OMVs in urosepsis

The mode of action of HlyF in the development of more severe UTIs remains elusive. HlyF induces the formation of OMVs [10]. OMVs can play the role of cargos. For example, it has recently been shown that UPEC OMVs can contain and transport the enzyme AroB, which is responsible for the synthesis of aromatic amino acids, thereby increasing the motility of recipient bacteria [45]. They can also increase the secretion of toxins such as CNF1 or CDT [10,46]. However, most of the *hlyF*+ UPEC strains, including ECC166 used in this study in the mouse UTI model, do not produce such toxins. It is therefore more likely that the intrinsic properties of OMVs could explain the pathogenicity of the strain producing HlyF. We have previously shown that OMVs produced by *hlyF+ E. coli* possess exacerbated pro-inflammatory properties that could play a role in the pathophysiology of infections and their severity [11].

### HlyF and pColV plasmid as virulence and antimicrobial resistance genes platforms: beyond UTI

HlyF displays wide dissemination in UPEC, suggesting that *hlyF*, and/or other determinants harboured by pColV plasmids, could give an advantage in a wide range of genetic backgrounds of *E. coli* for inducing UTI. However, we notice that major sequence types (ST) of *hlyF*+ strains also include NMEC and APEC strains which share an important genetic proximity. This is the case for ST95, one of the predominant UPEC ST in our collection and a highly represented ST within the *hlyF*+ UPEC strains [36,47,48]. However, other ST such as ST58, ST117 or ST131-*H22* are often strongly associated with avian infections which emergence and pathogenicity seem to be partly linked to the acquisition of a pColV plasmid [49–51].

*hlyF* and pColV plasmids are much more widely distributed than just in APEC, NMEC and UPEC and may contribute to the acquisition of virulence factors in other pathovars and to emergence of “high-risk” pathogenic clones [52]. Recently, a new emerging hybrid clone and lineage of *E. coli* has been described: an entero-haemorrhagic (EHEC) of serotype O80:H2 possessing typical EHEC virulence factors, which has acquired a pColV-like plasmid carrying *hlyF* and virulence factors usually associated with extra-intestinal pathogenic *E. coli* ExPECs [53,54]. Acquisition of this plasmid seems to give the EHEC host an atypical pathogenicity with a greater propensity to generate bloodstream infections, a known characteristic of ExPEC. Beside carrying the genes coding for various virulence factors, these plasmids are becoming concerning vectors of antimicrobial resistance genes, as seen for EHEC O80:H2 [55]. Other examples include the emerging ExPEC ST58 pColV+ sublineage and adherent-invasive *E. coli* strain NRG857c associated to Crohn disease, as well as an epidemic clone of *Salmonella enterica* serovar Kentucky: in both cases, the bacterial hosts harbour pColV-related plasmids that carry antimicrobial resistance determinants [49,54,56,57]. While these plasmids are all derived from the same conserved structure, their additional genes coding for antimicrobial resistance mechanisms may favour their selection and dissemination.

In conclusion, the results of our study and the sequencing of increasing numbers of *E. coli* isolates reflect a paradigm shift in our understanding of pColV plasmids: from being confined to APEC and NMEC, to their dissemination within new STs or even new *E. coli* pathotypes. Given the ageing of the population and the increase in co-morbidities and complex medical procedures such as the use of transplants or immunosuppressive drugs, the frequency and severity of UTIs is likely to increase [58]. Moreover, antimicrobial resistance is set to become one of the leading causes of mortality in the coming decades. It is therefore essential to monitor in clinical isolates the presence and evolution of plasmids that promote the dissemination of virulence factors such as HlyF, favour the onset of bloodstream infections, and carry antibiotic resistance genes.

## Acknowledgments

We thank the staff of the Tri GenoToul imaging facility, Toulouse.

## Data availability statement

Sequencing data (Illumina and Nanopore for some strains) are available in the NCBI Database, Bioproject number PRJNA615384.

## Authors contribution statement

Conceptualization: CVC, JPN, EO

Data curation: CVC, CM

Formal analysis: CVC, DP

Funding acquisition: CVC, JPN, EO

Investigation: CVC, DP, AG, CG, LD, MO, PJB, CS, CM, PB, JPN, MM

Methodology: YO, DP, LD, PB, JPN, MM

Project administration: EO

Resources: AG, PJB, CM, PB, MM

Visualization: CVC, DP, MO

Writing –original draft: CVC, DP

Writing –review & editing: CVC, JPN, MM, EO

---

**Supplementary Figure S1. *hlyF* is widely disseminated in UPEC strains**. A phylogenetic tree based on whole genome analysis of *hlyF*+ strains was constructed with *E. coli* MG1655 (*hlyF*-) as reference. Size of the circle is proportional to the number of *hlyF*+ strains of each ST, indicated in square brackets. Phylogroups are circled.

**Supplementary Figure S2**. Body weight gain/loss was compared before and at the end point of the experiment. Mean values ± SEM are shown, each circle represents a mouse. The presented results are pooled from two independent experiments. Student’s t-test: * p<0.05.

Supplementary Figure S3. Complementation restores the pro-pathogenic effect of HlyF during UTI.

C3H/HeN mice were infected trans-urethrally with ECC166 Δ*hlyF* transformed with a plasmidcarrying a wild-type *hlyF* gene (HlyF) or the *hlyF* gene with mutation in the catalytic domain (SDM).

**A.** Time to humane euthanasia (upon a body weight loss and/or clinical score reaching a predefined threshold) was monitored to build the survival curve. The difference between the experimental groups was evaluated by the log-rank (Mantel-Cox) test.

**B.** Clinical score according to Table S1 at 20h+/-2h post-inoculation.

**C.** Bacterial load in spleen (CFU/g organ) at end point.

**D.** Inflammatory cytokines in spleen (pg/mg proteins) at endpoint.

For B, C, D, mean values ± SEM are shown. Each circle represents a mouse. Student’s t-test: * p<0.05; ** p<0.01.

**Supplementary Figure S4. Global structure of ECC166 pColV conjugative plasmid carrying *hlyF*.** Genes of interest (virulence, plasmid replication, antibiotic resistance, conjugation system) are marked and coloured.

**Supplementary Figure S5. Co-phylogenetic relationship between *hlyF* and *traM-traX* locus.** The links between *hlyF* (left) and *traM-traX* (right) are indicated by grey lines. Strain names are color-coded according to their phylogenetic clusters determined by using RAMI.

