## Supplementary figures and images for "HlyF, an underestimated virulence factor of uropathogenic *Escherichia coli*"

### Figure S1

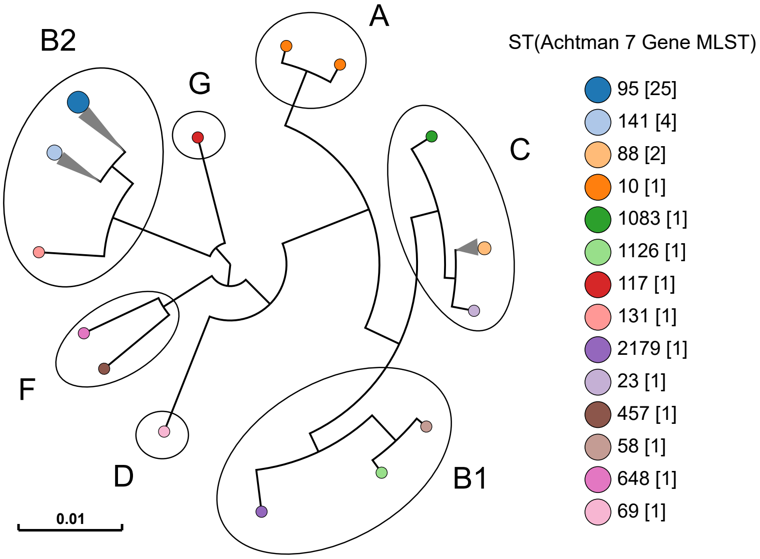

### Figure S2

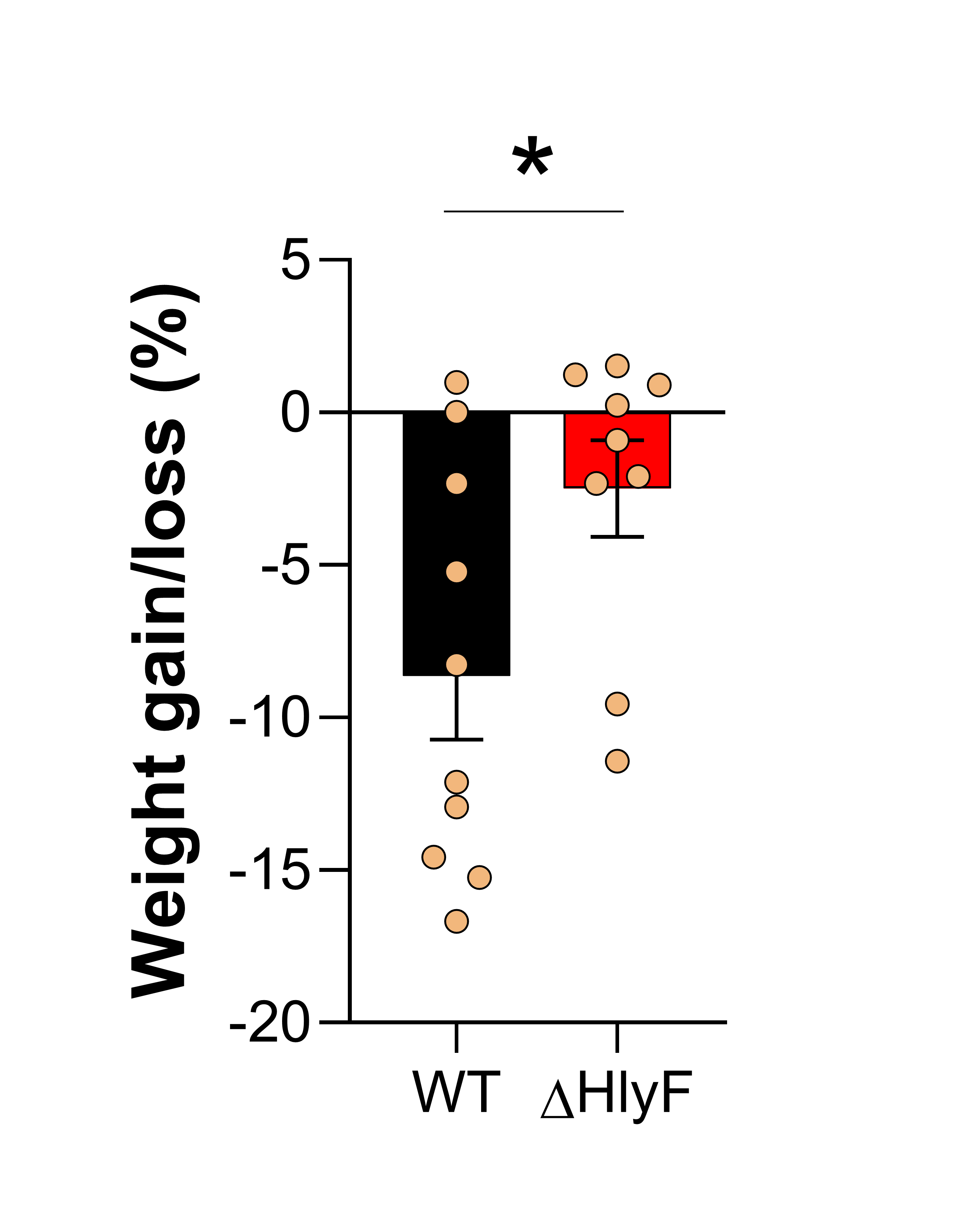

### Figure S3

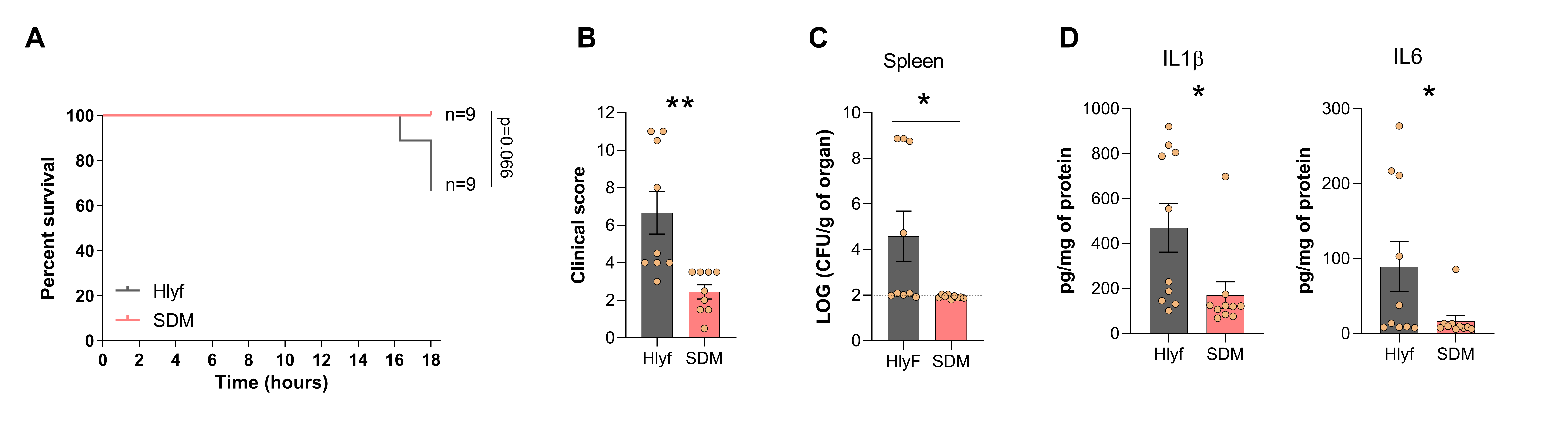

### Figure S4

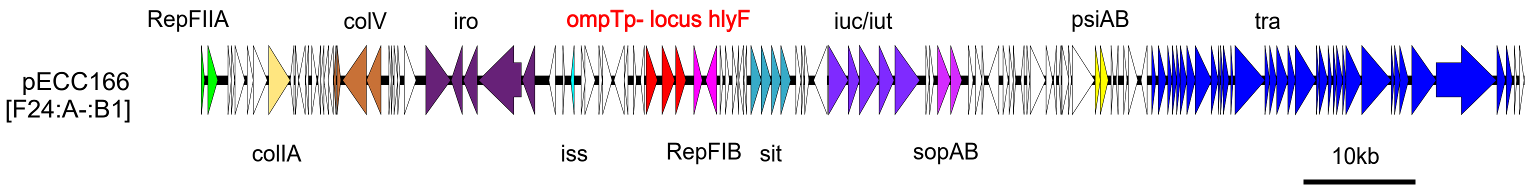

### Figure S5

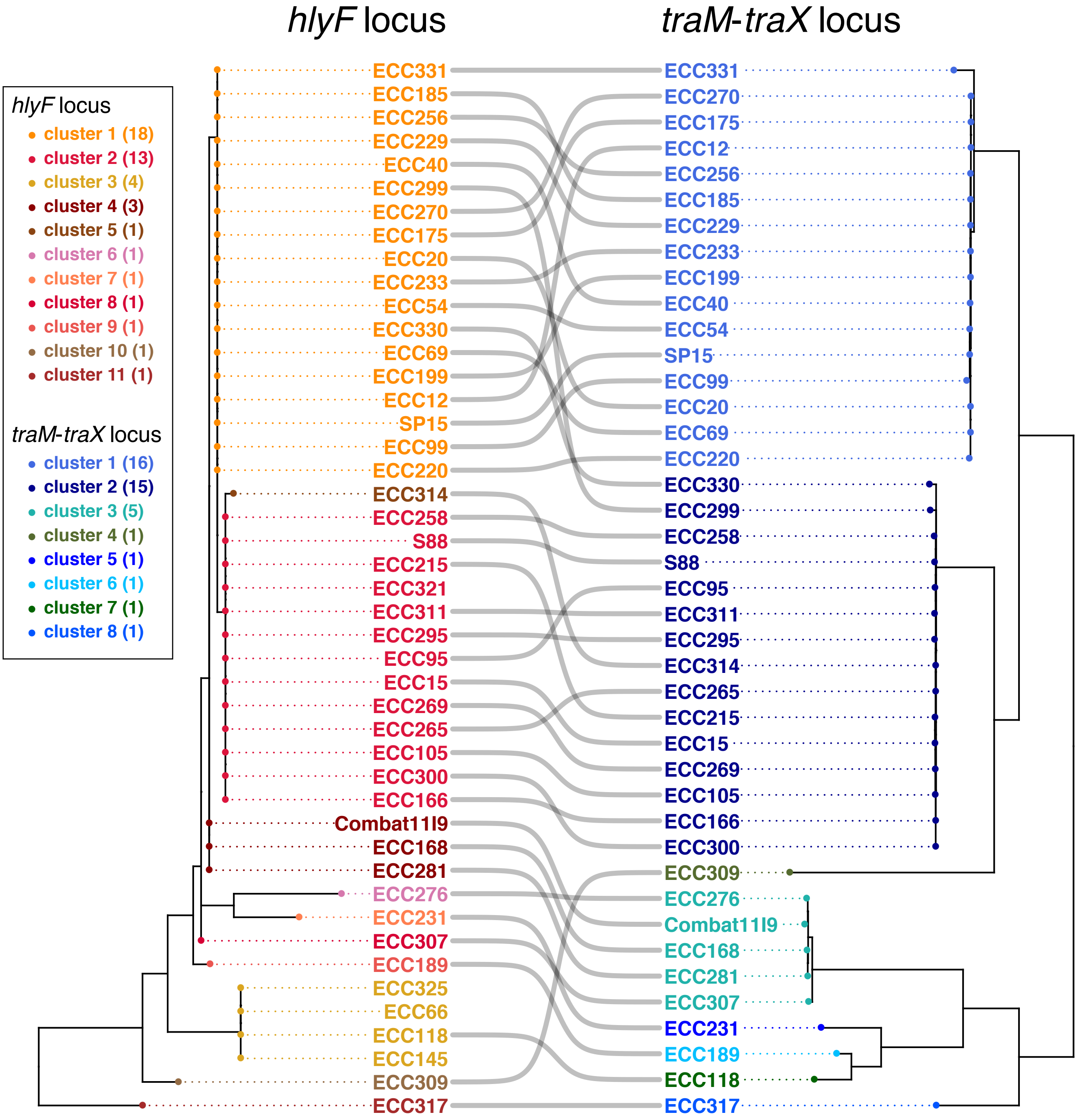
