## Supplementary material for "HlyF, an underestimated virulence factor of uropathogenic *Escherichia coli*": Table S2

**Supplementary Table S2:** SNP in *hlyF* locus compared to *E. coli* S88 reference strain (CU928146.1: 130 790… 133 302). Colours highlight phylogroups following the same legend as in the Figure 5.

|  | Position reference | Strain | NA change | AA change |
| --- | --- | --- | --- | --- |
| Promoter region | |  |  |  |
|  | 7 | 317 | g->a |  |
|  | 23 | 317 | c->t |  |
|  | 84 | 317 | g->t |  |
|  | 106 | 317 | g->a |  |
|  | 116 | 317 | t->g |  |
|  | 118 | 317 | a->c |  |
|  | 156 | 317 | a->c |  |
|  | 165 | 66;118;145;325 | c->t |  |
|  | 216 | 231;276 | g->t |  |
|  | 231 | 66;118;145;325 | c->t |  |
|  | 240 | 66;118;145;325 | c->t |  |
|  | 252 | 317 | t->g |  |
|  | 262 | 317 | g->a |  |
|  | 274 | 317 | a->c |  |
|  | 277 | 231 | a->g |  |
|  | 278 | 317 | c->a |  |
|  | 294 | 317 | c->t |  |
|  | 295 | 66;118;145;325 | a->t |  |
|  | 304 | 309 | c->t |  |
|  | 314 | 66;118;145;309; 317 ;325 | t->c |  |
|  | 318 | 66;118;145;325 | c->t |  |
|  | 320 | 317 | g->a |  |
|  | 329 | 317 | t->c |  |
|  | 352 | 309 | c->a |  |
|  | 372 | 317 | c->a |  |
|  | 375 | 66;118;145;325 | t->a |  |
|  | 376 | 309 | g->a |  |
|  | 408 | 66;118;145;309; 317 ;325 | c->t |  |
|  | 410 | 309 | c->t |  |
|  | 429 | 317 | a->g |  |
| hlyF: from 433 to 1541 | |  |  |  |
|  | 441 | 66;118;145; 231 ;309; 317 ;325; | a->g | silent |
|  | 688 | 314 | g->a | D85N |
|  | 759 | 189 | g->t | silent |
|  | 1096 | 276 | c->a | L221M |
|  | 1119 | 276 | c->t | silent |
|  | 1164 | 66;118;145;325 | c->t | silent |
|  | 1265 | 309;317 | a->t | D277V |
|  | 1286 | 317 | c->t | T284I |
|  | 1326 | 309;317 | t->c | silent |
|  | 1410 | 231 | t->c | silent |
|  | 1453 | 231 | c->t | P340S |
| intergenic |  |  |  |  |
|  | 1543 | 276 | t->a |  |
|  | 1572 | 276 | g->t |  |
|  | 1585 | 66;118;145;325 | t->c |  |
|  | 1586 | 317 | a->g |  |
| mig14: from 1605 to 2513 | |  |  |  |
|  | 1607 | 309;317 | g->a | M1I |
|  | 1637 | 276 | t->a | silent |
|  | 1814 | 276 | c->t | silent |
|  | 1859 | 231;276 | t->c | silent |
|  | 1889 | 231 | c->t | silent |
|  | 1913 | 231 | g->t | silent |
|  | 1922 | 231;276 | t->c | silent |
|  | 1947 | 276 | c->g | Q114A |
|  | 1948 | 276 | a->c | Q114A |
|  | 1985 | 231;276 | a->g | silent |
|  | 2039 | 231 | c->t | silent |
|  | 2055 | 66;118;145;168;189;231;276;281;307;  309;317; 325; pSP15 | c->a | silent |
|  | 2076 | 276 | g->a | E157K |
|  | 2094 | 12;20;40;54; 66;69;99; 118;145;168;175;185;  189;199;220;229;231;233;256;  270;276;281;299;307;309;  317; 325;330;331 ;pSP15 | a->c | K163Q |
|  | 2147 | 189 | g->a | silent |
|  | 2159 | 276 | c->g | silent |
|  | 2342 | 66;118;145;189;309 ; 325 | c->t | silent |
|  | 2393 | 66;118;145;189;231;307; 309; 317 ;325 | g->a | silent |
